# Adherence to the iDSI reference case among published cost-per-DALY averted studies

**DOI:** 10.1101/432377

**Authors:** Joanna Emerson, Ari Panzer, Joshua T. Cohen, Kalipso Chalkidou, Yot Teerawattananon, Mark Sculpher, Thomas Wilkinson, Damian Walker, Peter J. Neumann, David D. Kim

**Affiliations:** Center for the Evaluation of Value and Risk in Health, Tufts Medical Center, Boston, MA; Institute of Global Health Innovation, Imperial College London, London, UK; The Saw Swee Hock School of Public Health, National University of Singapore, Singapore; Centre for Health Economics, University of York, York, UK; Health Economics Unit, School of Public Health and Family Medicine, University of Cape Town, South Africa; Bill & Melinda Gates Foundation, Seattle, WA

## Abstract

**Background:** The iDSI reference case, originally published in 2014, aims to improve the quality and comparability of cost-effectiveness analyses (CEAs). This study assesses whether the development of the guideline has improved the reporting and methodology for CEAs using disability-adjusted life-years (DALYs).

**Methods:** We analyzed the Tufts Medical Center Global Health CEA Registry to identify cost-per-DALY averted studies published from 2011 to 2017. Among each of 11 principles in the iDSI reference case, we translated all reporting standards and methodological specifications into quantifiable yes/no questions and awarded articles one point for each item satisfied. We then separately calculated reporting and methods scores, measured as percent adherence (0%=no adherence, 100%=full adherence). Using the year 2014 as the dissemination period, we conducted a pre-post analysis. Additionally, we conducted an analysis stratified by the 11 principles and examined different scoring strategies and dissemination periods in sensitivity analyses.

**Results:** Articles averaged 74% adherence to reporting standards and 60% adherence to methodological specifications. Adherence to reporting standards increased slightly over time (72% pre-2014 vs. 75% post-2014, p<0.01), but methodological adherence did not significantly improve (59% pre-2014 vs. 60% post-2014, p=0.53). Overall, reporting adherence scores exceeded methodology adherence scores (74% vs. 60%, p<0.001). Articles seldom addressed budget impact (9% reporting, 10% methodology) or equity (7% reporting, 7% methodology).

**Conclusions:** The iDSI reference case has substantial potential to serve as a useful resource for researchers and policy-makers in global health settings, but greater effort to promote adherence and awareness is needed to achieve its potential.

## Background

Since the original Panel on Cost-Effectiveness in Health and Medicine proposed the use of a reference case as a benchmark of quality and methodological rigor (1, 2), various guidelines for conducting economic analyses have been proposed (3, 4). Over the last two decades, many countries, particularly high-income ones, have developed their own reference cases to inform decision-making in their health care systems (5–8). In contrast, most low- and middle-income countries (LMICs) have not developed such guidelines, possibly due to their limited capacity to do so (9).

To address the need for a reference case that could apply to different contexts, particularly in LMICs, the Bill and Melinda Gates Foundation (BMGF) supported the development of the Gates Reference Case for Economic Evaluation to ensure high quality and transparent analyses (10). The first version was published in 2014 as the Gates Reference Case and, later in 2016, was renamed the International Decision Support Initiative (iDSI) Reference Case (10, 11) to convey the breadth of its intended applicability. The iDSI Reference Case fills a major gap in global health economics, as it serves as the only resource of best practices for economic evaluation for many LMICs looking for guidance on resource prioritization. To date, however, no study has examined the extent to which economic evaluations adhere to the iDSI guidelines. We aimed to evaluate whether the development of the iDSI reference case has improved adherence to best practices for economic evaluations in global health settings, particularly cost-effectiveness analyses (CEAs) using disability-adjusted life years (DALYs).

## Methods

### Data

#### The iDSI Reference Case

The iDSI reference case includes 11 principles: transparency, comparator, evidence, measures of health outcome, costs, time horizon/discount rate, perspective, heterogeneity, uncertainty, budget impact, and equity considerations. Each principle has a number of corresponding methodological specifications to guide study design, and reporting standards to inform the communication of findings (Table 1). By using this tiered structure, the Reference Case aims to serve as a framework that both provides best practice guidance while allowing for flexibility depending on context. (11)

**Table 1:** The iDSI reference case: simplified principles, methodological specifications and reporting standards.

| **Reference Case Principle** | **Methodological specification, simplified** | **Reporting standards, simplified** |
| --- | --- | --- |
| Transparency | Decision problem, limitations, and declarations of interest are appropriately characterized. | Decision problem (population, intervention, comparator, outcome), evaluation's limitations, and declarations of interest are fully described. |
| Comparator(s) | Intervention(s) currently offered to the population (standard of care) is the base case comparator. | Comparator and its availability is clearly stated, and outcomes reported in incremental cost effectiveness ratio. |
| Evidence | Systematic literature review is used as source of evidence. | Methods of evidence collection are stated and sources of parameters are cited. |
| Measure of health outcome<sup>†</sup> | DALYs are used as the base case outcome measure. | Methods for weighting of DALYs are stated. |
| Costs | Costs are relevant to the context and stated perspective, and include implementation costs. | Costs are reported in local currency and USD. |
| Time horizon and discount rate | Lifetime time horizon and 3% discount rate for costs and outcomes are used in base case. | Time horizon and discount rate are clearly stated. |
| Non-health effects and costs outside health budget (perspective) | Societal perspective is used in base case, and relevant costs to this perspective (including direct health costs) are included. | Perspective and base case outcomes are clearly stated. |
| Heterogeneity | Heterogeneity is analyzed for appropriate subgroups. | Subgroup characteristics and analysis of heterogeneity are clearly described. |
| Uncertainty | Sensitivity analyses are performed on parameter source uncertainty (deterministic), parameter precision (probabilistic), and analysis structure (structural). | Magnitude of uncertainty in the model's structure, parameters, and precision are reported. |
| Budget impact | Intervention(s) budget impact is assessed. | Intervention(s) budget impact is reported. |
| Equity considerations | Intervention(s) implications on equity are assessed. | Intervention(s) implications on equity are stated. |

DALY: disability-adjusted life year; USD: United States dollar.
<sup>†</sup> The initial Gates reference case specified DALYs as a measure of health outcomes, but the 2016 iDSI Reference Case endorsed both QALY and DALY as appropriate measures because the focus should be placed on the principle to use outcome measures that are generalizable across disease states and capture positive and negative effects on both mortality and morbidity. Because all of our included studies used DALY as a measure of health outcomes, the change made in the 2016 iDSI Reference Case would not influence our results.

##### Global Health CEA Registry

We analyzed data from the Tufts Medical Center Global Health CEA Registry, a continually updated database of English-language economic evaluations in the form of cost-per-DALYs averted (12). Among 620 cost-per-DALY averted studies in the database, we selected a subset (N = 398) published three years before and after the initial release of the iDSI reference case (2011–2017) to examine the impact of its publication on the literature. We focused particularly on economic evaluations using the DALY metric because it is recommended as a main outcome metric by the iDSI Reference Case and it is used more often as a health outcome measure in LMICs than equivalent metrics such as the Quality-Adjusted Life Year (QALY). (11, 13)

To ensure a comprehensive assessment of adherence to the reference case, two independent readers (JE and AP) extracted additional information from each study in our sample using REDCap, an online survey platform (14), including data on: currency reported; subgroup analyses conducted; limitations reported; structural sensitivity analyses conducted; budget impact conducted; justification of alternative methodology; and comparator setting.

#### Adherence score

We first translated all 30 methodological specifications and 38 reporting standards (across 11 principles) listed in the reference case into questions with discrete yes/no outcomes (Appendix S1). We then designated reference case elements as “required” or “optional” based on our interpretation of the language in the report. We deemed 19 methodological and 21 reporting specifications “required”.

## Appendix S1: Reference case evaluation method

Our base-case analysis examined adherence scores consisting only of “required” elements. We evaluated each published cost-per-DALY averted study’s adherence to reporting standards (0–21 items) and methodological specifications (0–19 items). We then recorded for each article an overall reporting adherence score (proportion of 21 reporting standards adhered to) and an overall methodological adherence score (proportion of 19 methodological specifications adhered to).

## Analysis

### Descriptive analysis

We examined the association between adherence score and certain study characteristics, including whether the study cited the reference case, the study funder characteristics, and journal attributes. We categorized study funders into the following groups (not mutually exclusive): academic, government, healthcare organization, industry, intergovernmental organization, BMGF, non-BMGF, and other. We also stratified selected articles into clinical versus non-clinical journals using SCImago Journal Rank’s subject categorization (medicine vs. health policy, public health, non-health) (15). Finally, we recorded 2016 journal impact factor quartiles and categorized studies as high impact (first quartile), medium impact (second quartile), or low-impact (third and fourth quartiles) (15).

### Statistical analysis

To examine whether the iDSI guideline has since its release in 2014 improved the reporting and methodology for cost-per-DALY averted studies, we calculated mean adherence scores by year from 2011 to 2017. We conducted a pre-post analysis of improvement in methodological and reporting adherence using Student’s t-test. We considered the year 2014 to be the reference case’s dissemination period, and hence did not include articles published during that year in our pre-post analysis. We also compared the overall methodological specifications and reporting standards adherence scores, stratified by the 11 principles, using Student’s t-test.

### Sensitivity analysis

We conducted three sensitivity analyses. First, we included the “optional” specifications in the calculation of adherence scores for a random 10% subset of the articles to explore the impact of including optional items in the adherence score. Second, we removed 2015 from the post-evaluation period, limiting it to 2016–17, to examine the influence of alternative assumed dissemination period durations. Third, we used an alternative classification rule to score the one required adherence item pertaining to the comparator principle. To score the required comparator item “adherent”, our base case required the analysis to include as the comparator an intervention explicitly referred to as “standard of care”, a designation that can represent a range of possible interventions. Our sensitivity analysis scored any listed comparator other than “do-nothing” interventions as adherent. We designated “do-nothing” interventions as non-adherent to remain consistent with the principle that standard of care must at least be “minimal supportive care […] provided for that specification indication and patient group” in this sensitivity analysis (16).

## Results

### Descriptive statistics

Among 398 cost-per-DALY averted studies published from 2011–2017, 215 (54%) focused on LMICs and 263 (68%) targeted communicable diseases, such as diarrhea, HIV/AIDs, tuberculosis, and malaria (Table 2). Articles averaged 74% adherence to the reference case’s reporting standards and 60% adherence to the methodological specifications (Table 3). No article achieved full adherence to either the methodological specifications or the reporting standards.

**Table 2:**
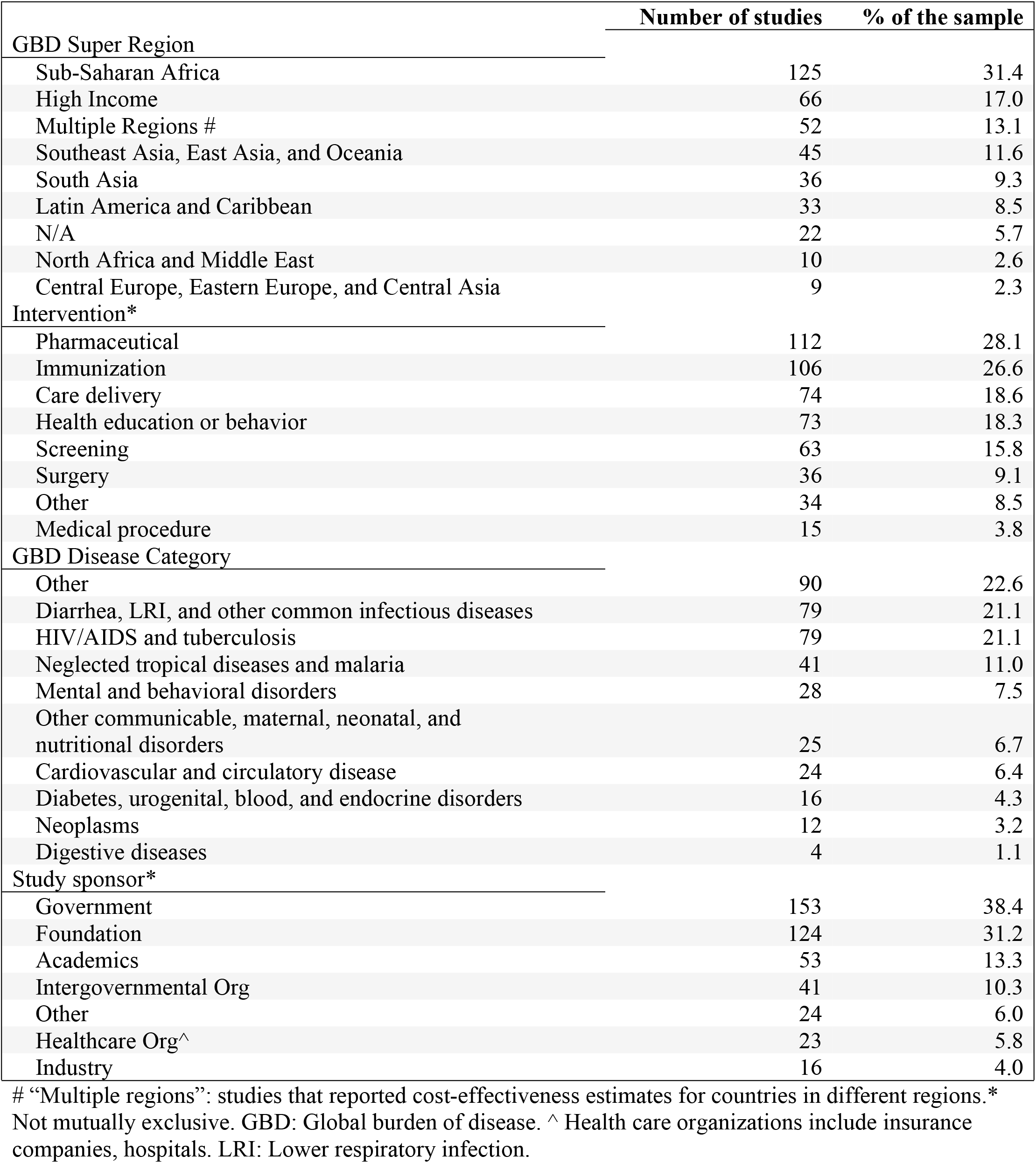
Characteristics of cost-per-DALY averted studies published 2011–2017 in Tufts Medical Center Global Health Cost-Effectiveness Registry.

|  | Number of studies | % of the sample |
| --- | --- | --- |
| **GBD Super Region** |  |  |
| Sub-Saharan Africa | 125 | 31.4 |
| High Income | 66 | 17.0 |
| Multiple Regions # | 52 | 13.1 |
| Southeast Asia, East Asia, and Oceania | 45 | 11.6 |
| South Asia | 36 | 9.3 |
| Latin America and Caribbean | 33 | 8.5 |
| N/A | 22 | 5.7 |
| North Africa and Middle East | 10 | 2.6 |
| Central Europe, Eastern Europe, and Central Asia | 9 | 2.3 |
| **Intervention\*** |  |  |
| Pharmaceutical | 112 | 28.1 |
| Immunization | 106 | 26.6 |
| Care delivery | 74 | 18.6 |
| Health education or behavior | 73 | 18.3 |
| Screening | 63 | 15.8 |
| Surgery | 36 | 9.1 |
| Other | 34 | 8.5 |
| Medical procedure | 15 | 3.8 |
| **GBD Disease Category** |  |  |
| Other | 90 | 22.6 |
| Diarrhea, LRI, and other common infectious diseases | 79 | 21.1 |
| HIV/AIDS and tuberculosis | 79 | 21.1 |
| Neglected tropical diseases and malaria | 41 | 11.0 |
| Mental and behavioral disorders | 28 | 7.5 |
| Other communicable, maternal, neonatal, and nutritional disorders | 25 | 6.7 |
| Cardiovascular and circulatory disease | 24 | 6.4 |
| Diabetes, urogenital, blood, and endocrine disorders | 16 | 4.3 |
| Neoplasms | 12 | 3.2 |
| Digestive diseases | 4 | 1.1 |
| **Study sponsor\*** |  |  |
| Government | 153 | 38.4 |
| Foundation | 124 | 31.2 |
| Academics | 53 | 13.3 |
| Intergovernmental Org | 41 | 10.3 |
| Other | 24 | 6.0 |
| Healthcare Org^ | 23 | 5.8 |
| Industry | 16 | 4.0 |

**Table 3:** Reference case adherence scores and percentages by year, sponsor, and journal aspects.

|  | N | Methodological specification score<br>(max 19) |  |  |  | Reporting standards score<br>(max 21) |  |  |  |
| --- | --- | --- | --- | --- | --- | --- | --- | --- | --- |
|  |  | Mean<br>(SD) | Min | Max | Mean<br>Adherence % | Mean<br>(SD) | Min | Max | Mean<br>Adherence % |
| Full sample | 398 | 11.3 (2.2) | 5 | 17 | 59.6 | 15.5 (1.8) | 9 | 19 | 73.9 |
| Year |  |  |  |  |  |  |  |  |  |
| Pre-2014 | 138 | 11.2 (2.3) | 5 | 17 | 58.9 | 15.2 (1.9) | 9 | 19 | 72.3 |
| 2014 and beyond | 260 | 11.4 (2.1) | 5 | 17 | 60.0 | 15.7 (1.7) | 10 | 19 | 74.7 |
| Study sponsor* |  |  |  |  |  |  |  |  |  |
| Academic | 53 | 11.3 (2.4) | 7 | 16 | 59.7 | 15.8 (1.7) | 11 | 19 | 75.0 |
| Government | 153 | 11.5 (2.1) | 6 | 17 | 60.7 | 15.8 (1.6) | 10 | 19 | 75.4 |
| Healthcare Org | 23 | 12.4 (2.1) | 6 | 17 | 65.4 | 16.2 (1.7) | 12 | 19 | 77.2 |
| Industry | 16 | 11.5 (2.1) | 6 | 14 | 60.5 | 15.7 (1.5) | 12 | 19 | 74.7 |
| Intergovernmental | 41 | 11.8 (2.0) | 7 | 16 | 61.9 | 15.6 (1.6) | 13 | 19 | 74.4 |
| Foundation | 56 | 11.4 (2.3) | 7 | 17 | 60.2 | 15.6 (1.7) | 12 | 19 | 74.1 |
| BMGF | 74 | 11.4 (2.1) | 6 | 16 | 60.1 | 15.8 (1.8) | 10 | 19 | 75.4 |
| Other | 24 | 11.1 (2.6) | 6 | 16 | 58.6 | 15.4 (1.9) | 11 | 19 | 73.2 |
| Cite reference case | 9 | 11.8 (2.4) | 9 | 17 | 62.0 | 16.6 (1.6) | 15 | 19 | 78.8 |
| Clinical journal | 318 | 11.5 (2.1) | 5 | 17 | 60.3 | 15.6 (1.7) | 10 | 19 | 74.2 |
| Non-clinical journal | 80 | 10.8 (2.5) | 5 | 15 | 56.8 | 15.2 (2.1) | 9 | 19 | 72.5 |
| Journal impact factor<sup>#</sup> |  |  |  |  |  |  |  |  |  |
| High | 336 | 11.5 (2.1) | 5 | 17 | 60.5 | 15.6 (1.8) | 9 | 19 | 74.1 |
| Medium | 45 | 10.7 (2.4) | 5 | 14 | 56.1 | 15.3 (1.6) | 12 | 19 | 73.0 |
| Low | 12 | 9.5 (1.3) | 8 | 12 | 50.0 | 14.8 (2.0) | 11 | 18 | 70.6 |

BMGF: Bill and Melinda Gates Foundation
\*Categories are not mutually exclusive. #Journal impact factor categories defined by 2016 SCImago Journal Rank quartile: high = first quartile; medium = second quartile; low = third and fourth quartiles. Five journals' impact factors were not available.

Of the 213 articles published after 2014 (i.e. 2015–2017), only 9 (4%) cited the iDSI reference case. For articles that did so, adherence to reporting standards averaged 79%, five percentage points higher than mean adherence for the full sample, while adherence to methodology specifications did not differ from adherence for the full sample. Funding source (BMGF vs. non-BMGF) was not significantly associated with adherence scores for either reporting (mean score of 75% vs. 74%) or methodology (mean score of 60% vs. 60%).

Studies published in clinical journals had marginally higher adherence (74% reporting adherence, 60% methodology adherence) than studies in non-clinical journals (73% reporting adherence, 57% methodology adherence). On average, methodology adherence scores for articles published in high-impact journals exceeded the corresponding scores for studies published in low-impact journals (61% vs. 50%); for reporting adherence, the corresponding difference was 74% vs. 71%.

Reporting standard adherence slightly increased after publication of the reference case compared to the pre-2014 period (72% adherence pre-2014 vs. 75% post-2014, p<0.01) (Figure 1). Methodological adherence did not improve (59% adherence pre-2014 vs. 60% post-2014, p = 0.53).

**Figure 1:**
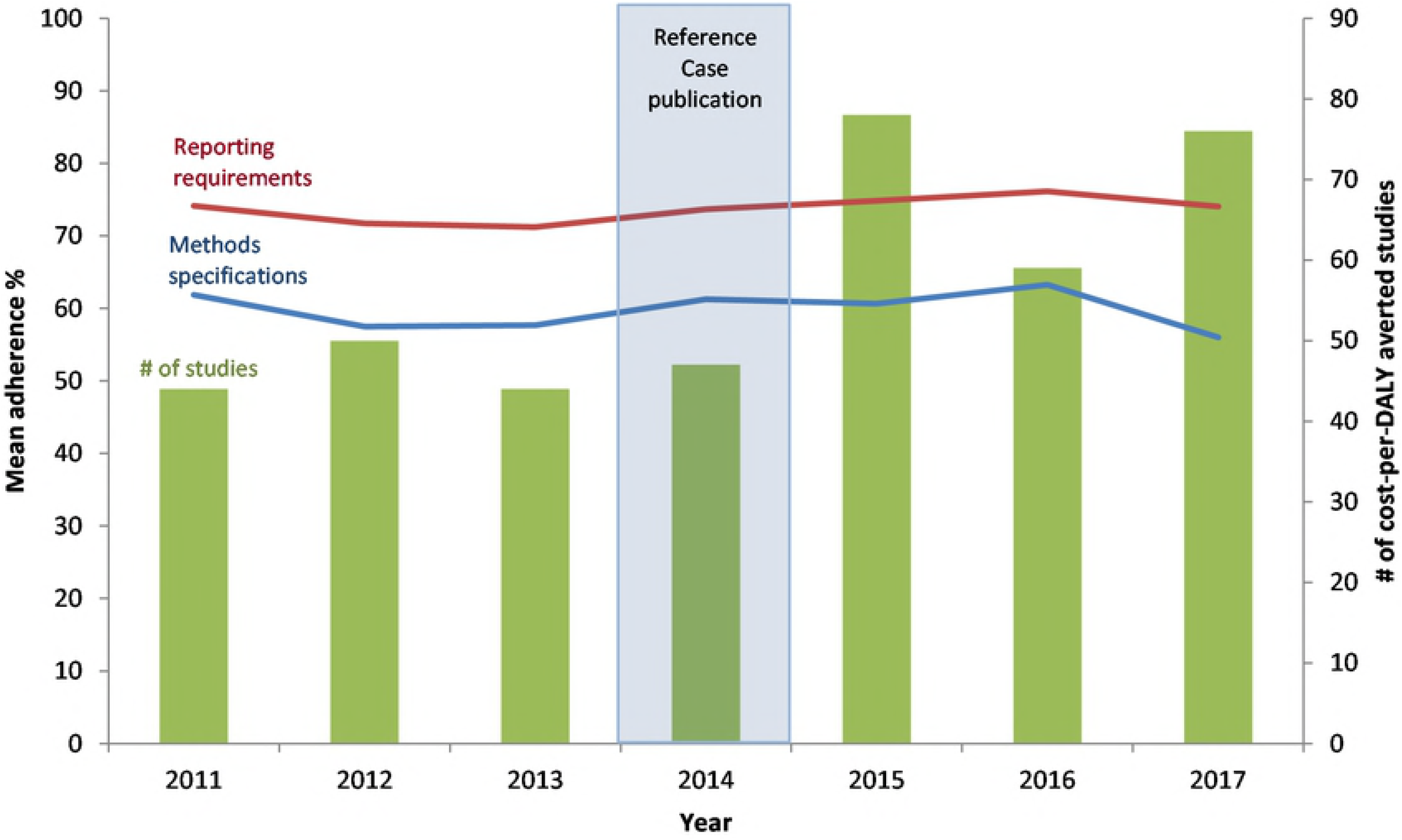
Reference case adherence percentages and number of cost-per-DALY averted studies over time.

### Methodological specifications versus reporting standards

Across the 11 principles, reporting standard adherence exceeded methodological specification adherence by 14 percentage points (74% vs. 60%). Reporting standard adherence was highest for the following principles: uncertainty (mean of 100%), comparator (97%), and evidence (95%). Methodological specification adherence was highest for the outcome measure (100%), transparency (89%), and evidence (74%) principles (Figure 2).

**Figure 2:**
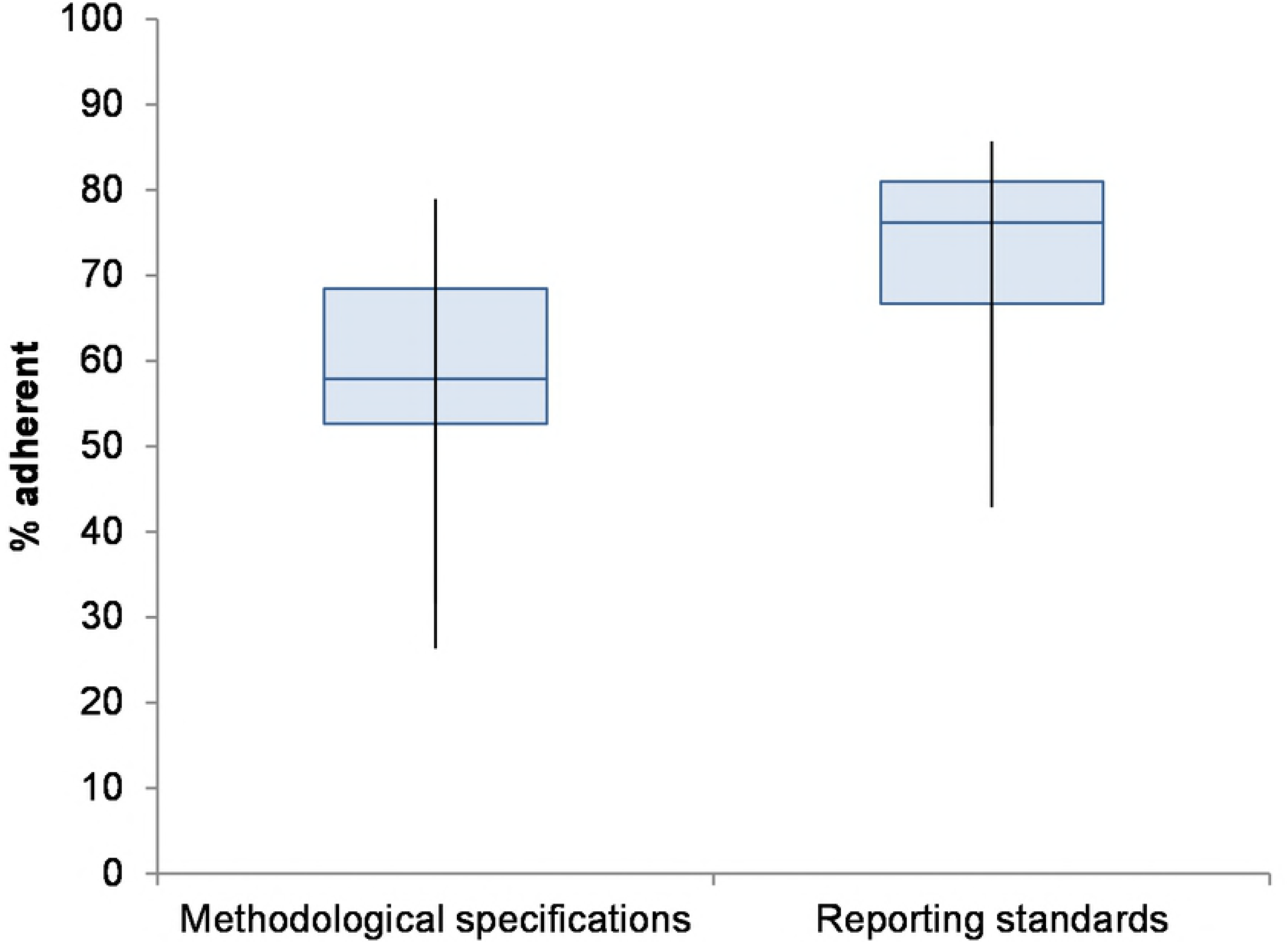
Box plot of article adherence percentage distribution for methodological specifications and reporting standards.

Reporting standard adherence exceeded methodological specification adherence for the following principles: comparator (97% vs. 36%), evidence (95% vs. 74%), time horizon/discounting (82% vs. 57%), perspective (85% vs. 64%), and uncertainty (100% vs. 57%) (Figure 3). Methodology adherence scores were higher than reporting adherence scores for the following principles: transparency (86% vs. 89%), outcome (54% vs. 100%), and costs (54% vs. 65%). Articles seldom addressed the budget impact (9% reporting adherence, 10% methodology) or equity (7% reporting adherence, 7% methodology) (Figure 3).

**Figure 3:**
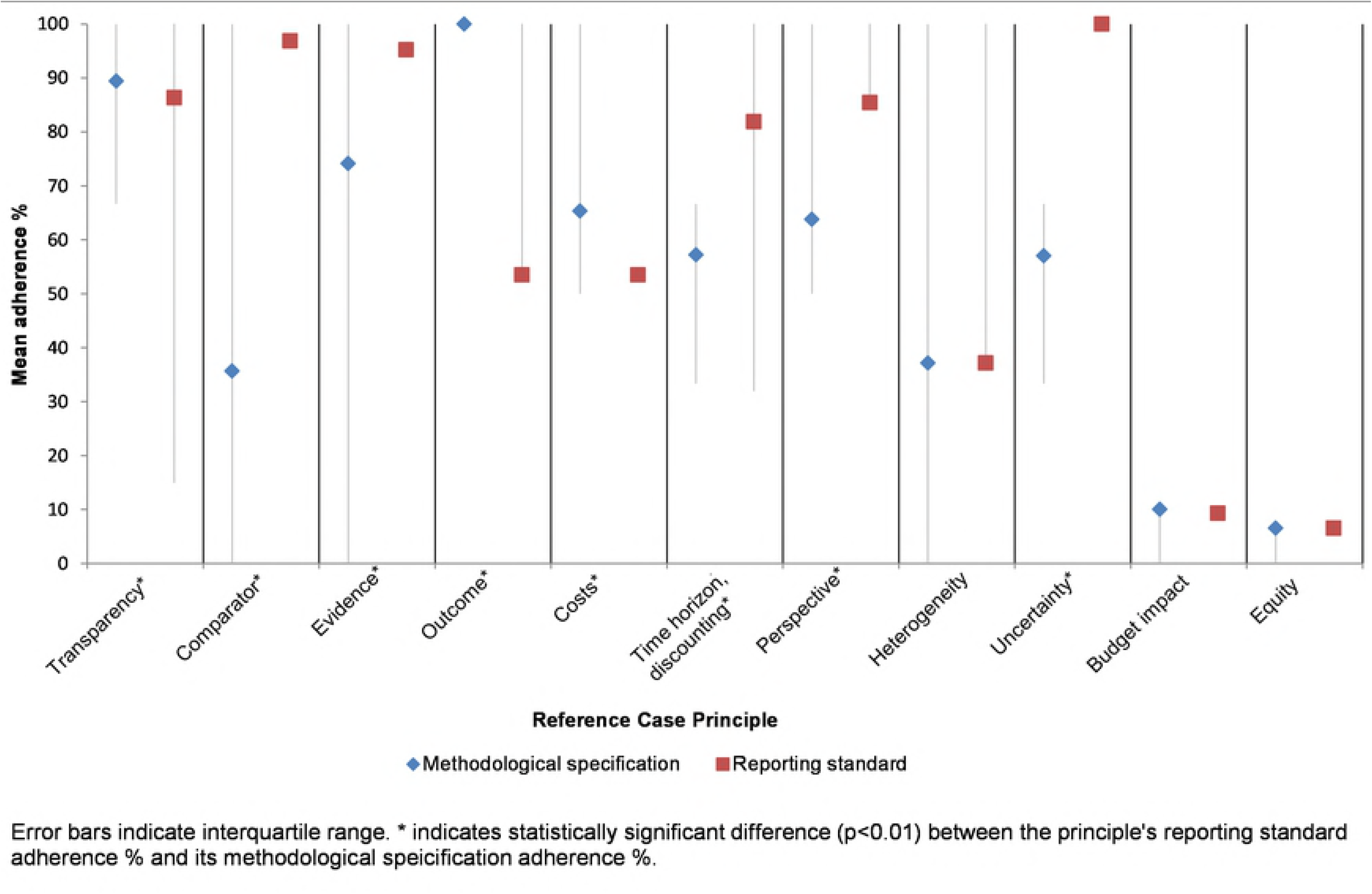
Percentage of articles adherent to reference case reporting standards compared to methodological specifications, by principle.

### Sensitivity analyses

Inclusion of optional criteria in our adherence score calculation decreased mean reporting adherence by 22 percentage points (from 74% to 52%) and mean methodological adherence by 14 percentage points (60% to 46%). When we limited the post-evaluation period to 2016–2017 (base case, 2015–2017), improvement in reporting standards post-publication no longer achieved significance. However, altering the comparator principle criteria (base case: comparator must be standard of care; alternative: comparator can be any intervention other than “do-nothing”) had little impact.

## Discussion

Since its release in 2014, adherence to the iDSI reference case among published cost-per-DALY averted studies has improved for reporting, but not for methods. We also found that adherence to the reference case’s reporting standards exceeds adherence to its methodological specifications, perhaps reflecting the relative ease of revising the way information is presented and greater effort needed to conform to analytic requirements. Moreover, other reporting guidelines, such as CHEERS (17), may have independently promoted more rigorous reporting, with the unintended effect of boosting adherence to the iDSI reporting standards

However, reporting and methodological adherence rates varied substantially across reference case principles, demonstrating ways in which articles are falling short of guidelines. For example, articles almost always report their comparator clearly (as recommended by the comparator reporting standard), but do not necessarily specify standard of care as the comparator (as recommended by the comparator methodological specification). Similarly, all articles reported findings from sensitivity analyses (as recommended by the uncertainty reporting standard), but did not always conduct structural, probabilistic, and deterministic analyses (as recommended by the uncertainty methodological specification). In some cases, methodological specification adherence exceeded reporting standards adherence. For example, articles often included implementation costs (as recommended by the costs methodological specification), but did not as frequently report these costs in both US dollars (USD) and local currency (as recommended by the costs reporting standards). Because the reporting and methods standards address distinct issues, future reference case specifications should continue to include both types of requirements.

The limited improvement in adherence to the iDSI reference case from 2011 to 2017 might reflect the competing influences of other guidelines, as authors may prioritize adherence to local guidelines or longstanding best practices (1, 8). The reference case aims to address the gap in guidance that exists for countries that cannot or have not yet created their own guidelines for economic evaluation. Local guidelines, which are tailored to their applicable context, may conflict with reference case guidelines; for example, the South African pharmacoeconomic guidelines recommend a 5% discount rate, which differs from the 3% value recommended by the reference case (18). Although the overall principle of the iDSI reference case supports the use of alternative discount rates where appropriate to the decision problem and constituency, researchers who adhere to the local guidelines may appear non-adherent to the methodological specifications as scored in this analysis.

Another possible explanation for relatively low adherence for certain items is that authors are not aware of the guidelines. Though the developers of the reference case have presented at various scientific meetings (19) and formally published the guidelines in 2016 (11), the BMGF and iDSI have focused educational campaigns on national payers and health technology assessment (HTA) agencies in LMICs, rather than on researchers, who are primary authors of published studies (16). Future studies should examine whether the reference case has influenced country-specific guidelines, such as Thailand’s HTA assessment guideline (6).

It is important to consider what level of adherence should be considered satisfactory. Although articles in our sample were more adherent to reporting guidelines, they adhered to just over half of methodological specifications. Adherence scores were notably lower for particular principles - heterogeneity, budget impact, and equity - indicating an overall neglect of these issues in cost-per-DALY averted studies. The adherence scores are perhaps best thought of as a baseline against which to measure improvement, and as a call to action to promote higher quality and comparability.

### Limitations

Our analysis has the following limitations. First, our use of dichotomous (i.e. “yes/no”) questions to score adherence may be inconsistent with the more nuanced goals of the iDSI reference case. Because the reference case is designed to be applicable in a range of different country-specific contexts, it must balance the goals of study comparability and quality against the goal of local applicability (16, 20, 21). To address this limitation, we omitted “optional” standards from our adherence calculation for the base case. That is, we assumed that the “optional” elements represent conditional requirements intended by the reference case authors to allow for local adaptability. Our sensitivity analysis that included all elements in our calculation of the adherence score (i.e. both the “required” and “optional” elements) yielded lower adherence scores.

Second, assessing adherence to the comparator methodological specification posed a particular challenge because this assessment depends on judging whether the specified comparator constitutes standard of care therapy. We explored the potential influence of our judgments by conducting a sensitivity analysis that redefined adherence to include any comparator other than “do nothing” interventions. This alternative itself posed a challenge because the “do nothing” intervention constitutes “standard of care” for some conditions in some settings. In any case, the fact that substantially altering this standard’s definition had little impact on our findings is reassuring.

Third, because the Tufts Medical Center Global Health CEA Registry catalogs only published cost-per-DALY averted studies, our findings cannot be generalized to the rest of the economic evaluation literature. For example, our analysis excluded gray literature (i.e., material not disseminated in regularly published, indexed journals). Gray literature may be more prevalent in some countries, especially those without local guidelines.

Fourth, our approach for scoring articles inherently involves reviewer judgement to determine author intent and to resolve ambiguities (e.g., determining whether the comparator is “clearly” stated). We attempted to mitigate this problem by having two reviewers read each article and, in cases where they could not reach agreement, appeal to a third reviewer.

Finally, our study’s post-evaluation period may not be sufficiently long to detect the impact of the reference case; as noted, the iDSI reference case was officially published in an academic journal in 2016 (11). More time may be needed for the field to adopt these guidelines.

### Policy implications

As posited by Nugent and Briggs, future research on the subject should ask, “what specific help does the iDSI reference case offer the analyst, who, while attempting to conform to the principles, nevertheless has to choose and implement the methods?” (22). It is possible that the methodological guidelines impose an excessive burden on researchers, raising “issues about the resources and data requirements to meet the principles” (16).

Future qualitative research should focus on researcher experience when conducting global health-focused CEAs and on how to increase its acceptance among authors. Studies could also evaluate the methods and reporting practices for articles that strongly adhere to the iDSI reference case, as these analyses may serve as useful examples for other CEA authors attempting to adhere to the guidelines. Future research should also evaluate the influence of the reference case on how decision makers perceive the quality and usefulness of economic evaluations.

Moving from guideline development to implementation is a vital step towards improving the utility of economic evaluations in global health. Future efforts could include additional educational workshops for researchers, students, and policymakers. Policymakers and major funders of economic evaluations, such as the BMGF, could require that researchers adhere to reference case recommendations.

## Conclusion

Our results indicate that the iDSI reference case has slightly improved reporting practices of economic evaluations focused on global health, but not methodological practices. The reference case has substantial potential to serve as a resource for researchers and policy makers in global health and economics, but more effort to promote adherence and awareness may be needed.

